# A kinetic ensemble of the Alzheimer’s Aβ peptide

**DOI:** 10.1101/2020.05.07.082818

**Authors:** Thomas Löhr, Kai Kohlhoff, Gabriella T. Heller, Carlo Camilloni, Michele Vendruscolo

## Abstract

The discovery that disordered proteins are widespread in the human proteome has prompted the quest for methods to characterize the conformational properties that determine their functional and dysfunctional behaviour. It has become customary to describe these proteins in terms of structural ensembles and free energy landscapes, which offer conformational and thermodynamic insight. However, a current major challenge is to generalize this description to ‘kinetic ensembles’, thereby also providing information on transition rates between states. Approaches based on the theory of stochastic processes can be particularly suitable for this purpose. Here, we develop a Markov state model and illustrate its application by determining a kinetic ensemble of the 42-residue form of the amyloid-β peptide (Aβ42), whose aggregation is associated with Alzheimer’s disease. Using the Google Compute Engine, we generated 315 μs all-atom, explicit solvent molecular dynamics trajectories, validated with experimental data from nuclear magnetic resonance spectroscopy. Using a probabilistic-based definition of conformational states in a neural network approach, we found that Aβ42 is characterized by inter-state transitions no longer than the microsecond timescale, exhibiting only fully unfolded or short-lived, partially-folded states. We contextualize our findings by performing additional simulations of the oxidized form of Aβ42. Our results illustrate how the use of kinetic ensembles offers an effective means to provide information about the structure, thermodynamics, and kinetics of disordered proteins towards an understanding of these ubiquitous biomolecules.

## Introduction

Proteins that are fully or partially disordered make up approximately one third of the human proteome, perform a variety of biological functions and are closely involved with many major human disorders^1–3^. The existence of disordered proteins is making it necessary to extend the structure-function relationship that has driven major advances in protein science in the last 50 years to a structure-dynamics-function relationship, in order to account for the essential role of structural disorder in determining the normal and aberrant behaviours of these proteins^1–4^.

Because of their conformational heterogeneity, it is typically insufficient to characterize disordered proteins using one or a few specific structures, as it is standard for folded proteins^5^. Instead, it has become common to describe these proteins in terms of structural ensembles, which in turn are often represented through free energy landscapes^6–9^, when the statistical weights of the states in the ensemble are available (for example, through the use of enhanced sampling techniques^6^). This is a powerful description, which concisely captures information about the structure and thermodynamics of disordered proteins. Here, we refer to these ensembles here as ‘thermodynamic ensembles’. It has also been recently observed, however, that a more complete description should include information about the kinetics, which can be achieved by adding the transition rates between different conformational states^10^. We refer to these ensembles here as ‘kinetic ensembles’. This task requires a characterisation of the kinetic properties of disordered proteins, which remains a challenging task, both experimentally and computationally.

In this work, we describe an approach to generate kinetic ensembles, which contain information about the molecular structures of proteins, the populations of their metastable states, and the transition rates between these different metastable states (**Figure 1a**). While acquiring structural and population information is already possible with molecular dynamics simulations alone, gaining interpretable kinetic information can be more effectively achieved by exploiting the theory of stochastic processes^11^ during the analysis, and it is typically done using Markov state models^12,13^.

**Figure 1.**
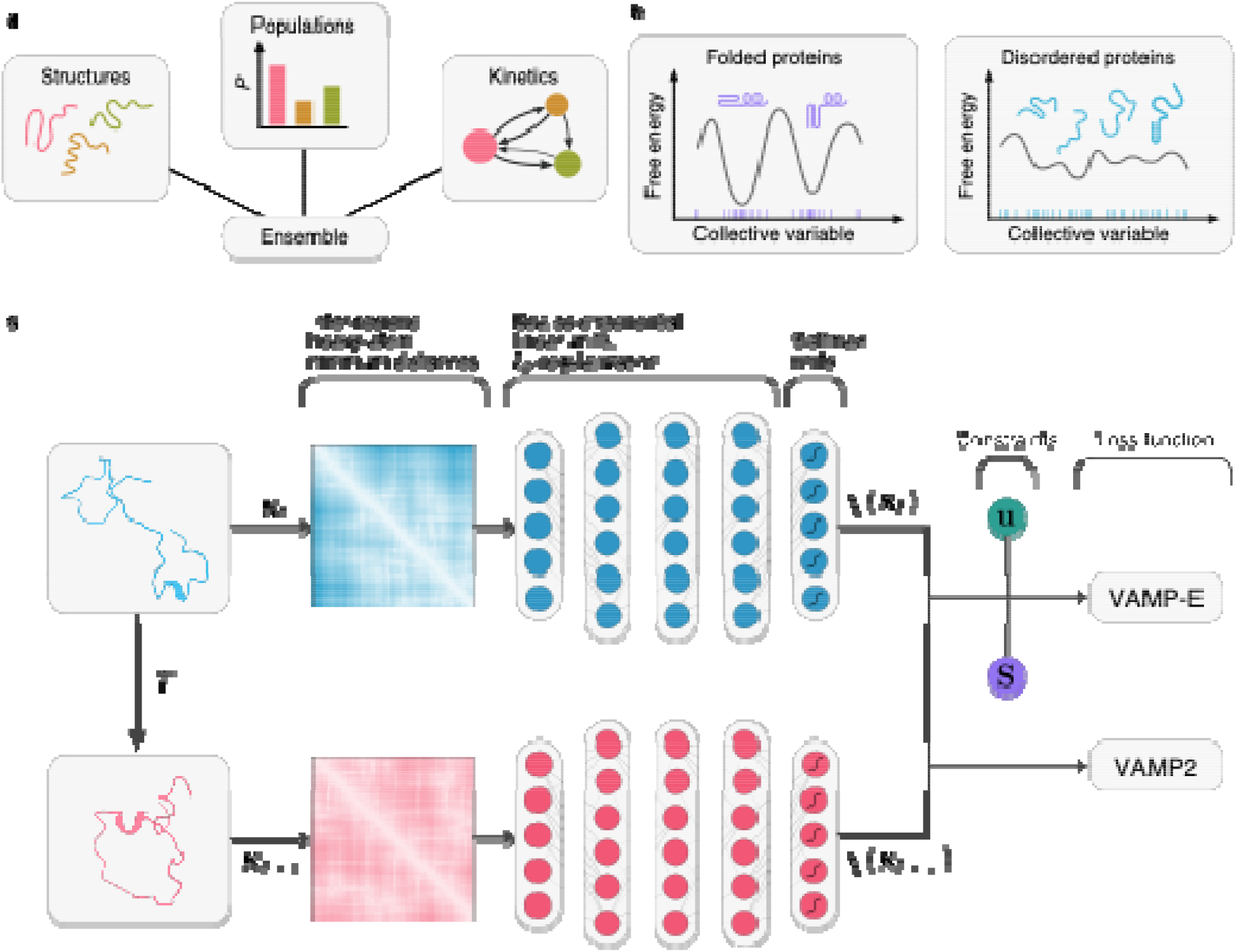
Schematic illustration of a kinetic ensemble of a protein and the training methodology. **(a)** A kinetic ensemble consists of three components: (1) structures, (2) their statistical weights (populations), and (3) interconversion rates between groups of related structures (states). Structural and thermodynamic ensembles have only the first or first two components, respectively. **(b)** The free energy landscape of disordered proteins is flatter and more heterogeneous than that of folded proteins, making a state decomposition difficult, and thus the determination of a kinetic ensemble challenging. **(c)** Architecture of the constrained VAMPNet. It consists of two lobes with shared weights, taking two frames separated by some lag time as input. The *χ* part is first trained with the VAMP2 score, followed by the constrained and full network with the VAMP-E score, respectively.

In contrast to thermodynamic ensembles, constructing kinetic ensembles requires a state decomposition. To define the states in these models, one can choose a set of features from a trajectory, such as backbone dihedral angles or the root-mean-square deviation to some reference structure, and then use a clustering algorithm to obtain a state assignment for each frame. One can then count the transitions between states and normalize these counts to obtain a transition matrix^12,14^. An additional coarse-graining step can then be performed to obtain a more interpretable model with fewer states. Alternatively, clustering can be preceded with an additional dimensionality reduction step. One such example particularly relevant for building Markov models is time-lagged independent component analysis (TICA)^15,16^ which projects conformations into a space where distances have a kinetic meaning. Clustering in this space thus has the potential of naturally preserving kinetic separation.

This particular model-building approach presents unique challenges for disordered proteins, due to the heterogeneous nature of their conformations. Transitions between metastable states are typically fast and not necessarily characterized by large variations in the overall shape of the protein. This is in contrast to folded proteins, where states can often be classified on structural properties alone^17–19^. For disordered proteins, even with suitable structural measures, because these transitions are fast, dividing this space into discrete areas is exceedingly difficult (**Figure 1b**), and many clustering algorithms may fail to consistently classify states. Consequently, it would be ideal to obtain a type of clustering, which we can describe as ‘soft’, such that state assignments can have a probabilistic nature. In this approach, every frame is assigned a discrete probability distribution encoding its membership in a certain state. This information can be used to build a soft Markov state model, which can be interpreted in much the same way as one built from individual transition counts^20^. To optimize for kinetic separation and allow for probabilistic state assignments, we employed the VAMPNet technique with physical constraints (**Figure 1c**)^21,22^ to construct a soft Markov state model (or Koopman model, see Methods). The resulting stochastic matrix allows us to compute the equilibrium distribution of the ensemble as well as the transition rates between metastable states.

We illustrate this approach to determine a kinetic ensemble of Aβ42, using 315 μs of explicit solvent molecular dynamics simulations. We chose Aβ42 because of its association with Alzheimer’s disease, which is among the leading causes of death in the developed world, with no effective treatments available^23^. One of the characteristics of Alzheimer’s disease is the formation of neurotoxic aggregates of Aβ42^24–29^. This complex process can be divided in to a set of nonlinearly coupled microscopic steps, for all of which the monomeric state of the protein plays a key role^30^. It is therefore of great importance to better understand the structural and kinetic properties of the monomeric state of Aβ42, as the dynamic properties in particular are key to determine the effects of small molecule drug candidates on the aggregation behaviour of this peptide^31,32^. Because of the ensemble-averaged nature of most experimental measurements of Aβ42, molecular simulations can provide a uniquely precise, atomistic description of its kinetic ensemble. Previous computational studies to characterize Aβ42^33–38^ have focused mainly on the thermodynamic properties of this peptide, but less so on the kinetics of state transitions. It is therefore still debated as to whether this peptide ever adopts long-lived (i.e. more than hundreds of microseconds) states associated with its aggregation behaviour. We validate the kinetic ensemble using independent experimental data obtained from nuclear magnetic resonance (NMR) spectroscopy. To the best of our knowledge, this is the longest published simulation of any disordered protein in explicit solvent executed to date. The resulting kinetic ensemble provides the structural properties and the populations of the states of Aβ42 and the transition rates between them. It also allows us to make predictions of involved timescales and is especially informative in terms of the kinetics of secondary structure formation. We believe that this approach offers a unique perspective into the structure, thermodynamics, and kinetics of disordered proteins towards an increased understanding of their dynamic behaviour.

## Results

### Generation of molecular dynamics simulations on the Google Compute Engine

We performed explicit solvent molecular dynamics simulations in 5 rounds with 1024 trajectories each, using the fluctuation amplification of specific traits (FAST) approach^39^ to accelerate the exploration of conformational space. Cloud architectures, such as the Google Compute Engine are especially well suited to generate Markov state models, as the individual simulations can be run independently from one another, with no communication between machines required^40^. We chose the CHARMM22* over the CHARMM36m force field due to the better agreement with macroscopic observables such as radius of gyration for the closely related Aβ40 peptide^41,42^.

### Standard Markov model techniques do not readily describe the Aβ42 ensemble

We first attempted to build discrete Markov state models using existing state-of-the-art techniques. We used time-lagged independent component analysis (TICA)^15,16^ as a preliminary dimensionality reduction step with inter-residue nearest-neighbour heavy atom distances, followed by clustering with various algorithms, different numbers of input dimensions, and varying amounts of clusters. We used the cross-validated generalized matrix Rayleigh coefficient technique^43^ to score the individual models (see Methods, **Figure S1**). Despite an extensive hyperparameter search, we were unable to construct a model with converging timescales (**Figure S2a**). Nevertheless, we used the best-performing hyperparameters, specifically the minibatch *k*-Means clustering algorithm using 16 dimensional input and 200 clusters to build discrete coarse-grained models using the hidden Markov state model^44^ (**Figures S2b-d**) and Perron cluster-cluster analysis approaches^45^ (**Figures S2e-f**). These models, however, also suffered from non-converging or overly fast timescales, suggesting the presence of significant problems in this particular approach for state-space discretization. We therefore decided to pursue an approach based on a probabilistic state description.

### Soft Markov state models describe the Aβ42 ensemble

We next attempted to build a neural network for the disordered ensemble of Aβ42 with soft state assignments. To this end we employed the VAMPNet technique with physical constraints^21,22^. VAMPNet is a two-lobed, unsupervised neural network, only taking as input two frames separated by a lag time *τ*, and yielding a soft state assignment vector *χ*(**x**_*t*_) for each frame **x**_*t*_ (**Figure 1c**). The loss function is given by the variational approach to Markov processes (VAMP) score^46^, allowing the learning of a state decomposition without explicit state labelling. The dimensionality reduction and clustering steps are thus performed in a single procedure, allowing for highly non-linear state membership functions to be learned. A recent addition^22^ to this approach allows the use of constraints to keep the elements in the learned transition matrix positive, and the model reversible. This is accomplished by training two auxiliary weights **u** and **s** and using the VAMP-E loss function. The transition matrix can then be learned by first training the unconstrained VAMPNet, followed by the constraint vectors and finally all trainable parameters together. We adopted this constrained framework but employed a self-normalising neural network architecture^47^ to improve the hyperparameter search (see Methods). As input, we used inter-residue nearest-neighbour heavy atom distances, resulting in an input dimension of 780, with a network lag time of 5 ns. We used 2 to 6 output states, finding that the 4-state model struck the best balance between interpretability, detail, and state assignment errors (**Figures S3, S4**). Outputs with increased numbers of states resulted in larger errors and a lack of interpretability.

### Computational validation of the Aβ42 kinetic ensemble

To construct a kinetic model, a suitable lag time τ between successive conformations along a trajectory needs to be chosen. This lag time should be small enough for the model to effectively resolve fast degrees of freedom, but large enough to not introduce a significant statistical error in the prediction of longer timescales. To choose this parameter and evaluate the general quality of the model it is helpful to visualize the longer timescales – a kinetic property of the model – as a function of the lag time itself. To this end, we assessed the dependence of the relaxation timescales, *t*_*i*_, (also called implied timescales in this approach, see Methods) on the model lag time *τ* (**Figure 2a**), while keeping the network lag time constant at τ = 5 ns. We observed that a model lag time *τ* = 12.5 ns can resolve longer timescales well. To further validate this choice for the model lag time, we used the Chapman-Kolmogorov test (**Figures S5-7**), a stringent measure of the predictive abilities of our model. To evaluate the quality of sampling in the context of the final model, we estimated Koopman operators with a limited number of trajectories (**Figure 2b**) from the existing full model. We would expect the relaxation timescales to accelerate and converge with the number of trajectories utilized as we improve the sampling of each state transition. The timescales indeed converge to within the error of the model. To understand how the choice of the number of states impacts the corresponding state assignments of individual frames, we illustrate the state decomposition as a tree (**Figure 2c**). States can be seen to be mostly consistent across different output sizes.

**Figure 2.**
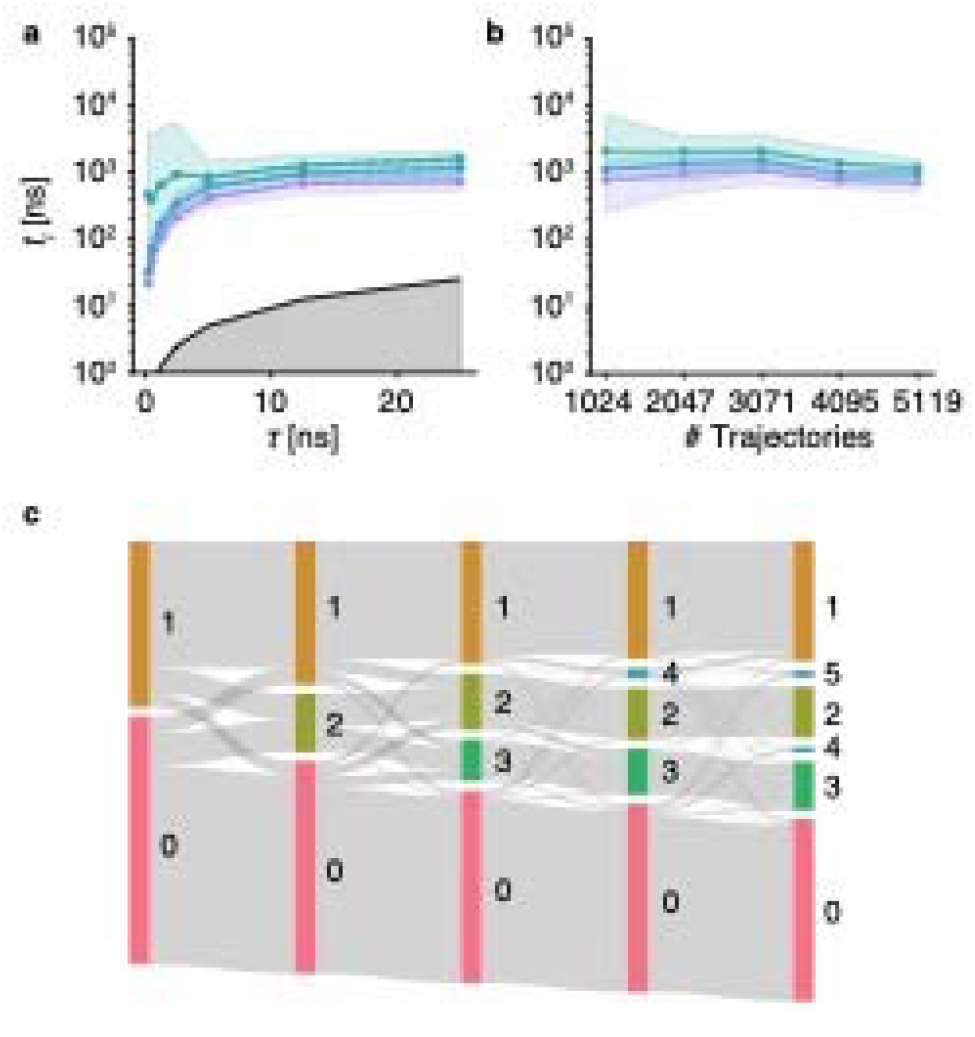
Determination of the states in the kinetic ensemble of Aβ42. **(a)** Dependence of the three longest relaxation timescales (green, cyan and purple, respectively) on the lag time *τ*. These timescales are derived from the eigenvalues of the transition matrix (see Methods). The grey shading indicates the timescales for which the Koopman model can no longer resolve the relaxation timescales. **(b)** Dependence of the relaxation timescales on the number of trajectories used to build the kinetic ensemble as a 4-state model. **(c)** Shift of equilibrium distributions and state assignments over different output sizes. Grey lines indicate state decomposition across multiple models with varying numbers of states, showing consistency across different output sizes. Shaded areas indicate 95th percentiles of the bootstrap sample of the mean.

### Experimental validation of the Aβ42 kinetic ensemble

We compared experimental NMR chemical shifts^31^ to values back-calculated from the simulations (**Figure S8a**). We found that the root-mean-square deviation between experiments and simulations is within the error of the forward model^48^. Additionally, we directly compared the distributions of the radius of gyration between the kinetic ensemble derived here and of a thermodynamic ensemble determined previously using metadynamic metainference^49,50^ simulations^31^, using the same force field and experimental restraints in the form of chemical shifts (**Figure S8b**). The relaxation times of around 2 μs (**Figure 2a**) are also consistent with experimental NMR measurements^51^. However, acquiring more precise data on state transitions of disordered proteins using NMR spectroscopy is limited by the vast structural variance of those proteins on short timescales and ensemble-averaged nature of the experiments^52^.

### Unconstrained model comparison

We compared the transition matrices, relaxation timescales and state lifetimes generated by constrained and unconstrained VAMPNets (**Figure S9**). We found that all timescales and transition probabilities are consistently slower in the unconstrained case. This is to be expected, as the estimation of the eigenfunctions is impeded by the constraints on non-negativity and reversibility^20^. However, as we are performing equilibrium simulations, we should expect detailed balance to hold. We therefore consider the constrained model to be better suited to this analysis. The difference can also be partially attributed to the use of many shorter trajectories, which introduces a bias in the unconstrained model^22^.

### Aβ42 states are structurally diverse

To characterize the local structural features and long-ranged self-interactions of the states in the kinetic ensemble, we calculated the α-helical and β-sheet content per residue (**Figures 3a, S10a** and **S11a**) and inter-residue contact probabilities (**Figures 3b, S10b** and **S11b**) in each state. We observed unique structural properties in each state and found that the C-terminal region of the peptide is generally more structured than the N-terminus. Specifically, each state is characterized by varying degrees of α-helix formation in localized areas, while the region between residues 10 and 20 generally remains completely disordered. The β-sheet content is also concentrated towards the C-terminal region with state 3 being a notable exception, partially forming an end-to-end contact. Moreover, many conformations identified by the secondary structure predictor DSSP as being rich in β-sheet content are characterized more by strong backbone interactions and less by explicit β-like structures, as can be seen in the contact probability maps. Using a cut-off of 0.45 nm we find an ensemble-wide proportion of 3.68 0.14 % of end-to-end contacts, comparable to the ensemble reported by Meng *et al*^33^.

**Figure 3.**
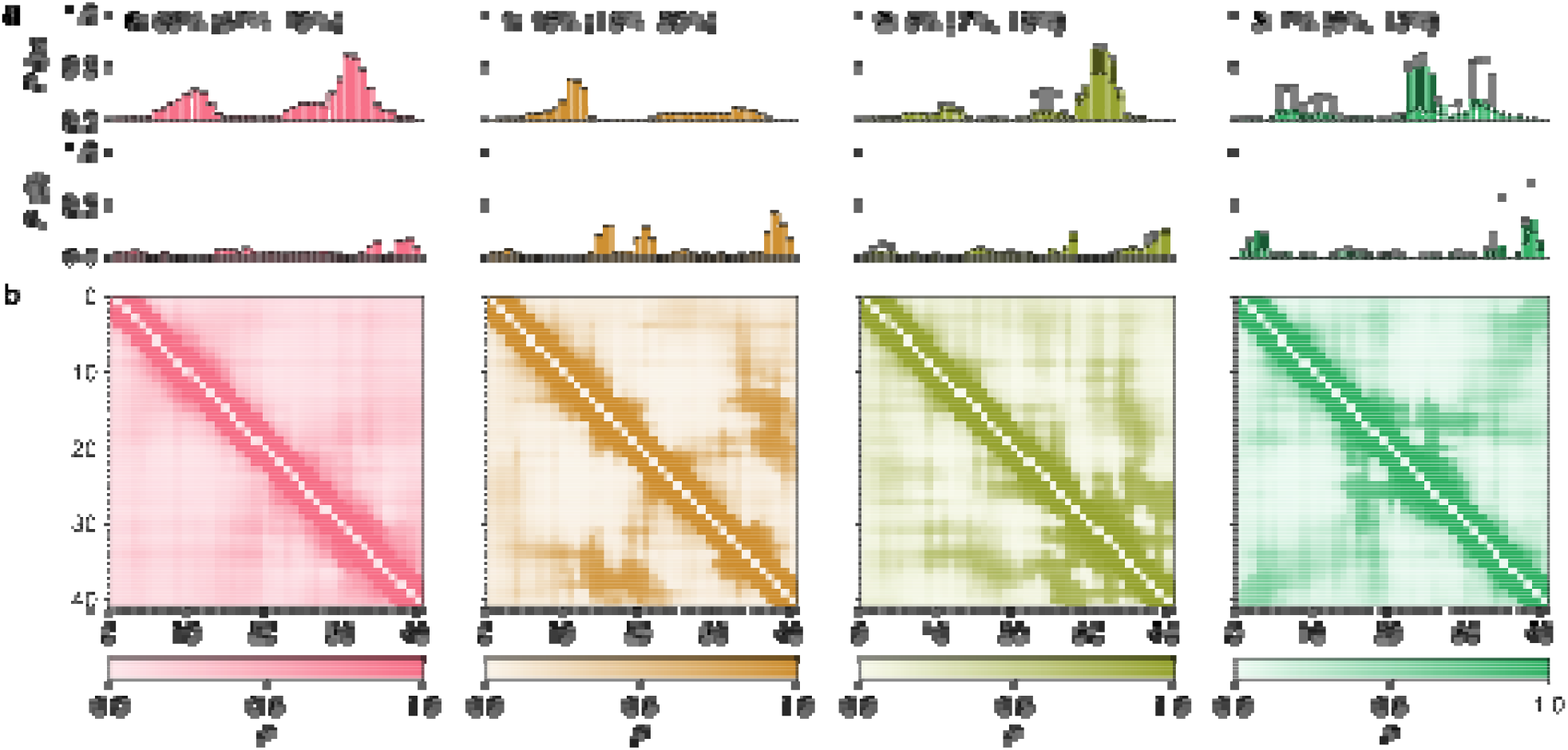
Structural properties of Aβ42 in the kinetic ensemble. **(a)** Populations of α-helical and β-sheet content per residue in each of the four states in the kinetic ensemble, as calculated using the secondary structure predictor DSSP. The equilibrium percentage of each state is given above, with the 95th percentile in parentheses. **(b)** Contact probability maps of each of the four states with a cut-off of 0.8 nm. Error bars indicate 95th percentiles of the bootstrap sample of the mean.

### Aβ42 is characterized by transitions between states on the μs timescale

The overall kinetic properties of a protein are primarily given by the inter-state transition probabilities (**Figure 4a**), the slowest relaxation timescales (**Figure 4b**), and mean state lifetimes (**Figure 4c**). We observe mean first-passage times between ~1 and ~12 μs, with no observation of longer-lived folded states. Instead, we observe the formation of a disordered hub-like state 0, to which other structured states transition relatively quickly. On the other hand, transitions into structured states, i.e. direct formation of secondary structural motifs, are significantly slower. Transitions bypassing the hub-state are rare, such as transitions between states 1 to 2. The relationship between the relatively fast relaxation times (~2 μs) (**Figure 4b**) and the somewhat slower mean first-passage times (1-12 μs) has previously been investigated for unfolded states of small proteins^53,54^. It was observed that the lifetime of a state is approximately equal to the timescale of relaxation to equilibrium (**Figure 4c**), thereby suggesting that the mean first-passage time into a state is mostly dependent on the population of the state itself.

**Figure 4.**
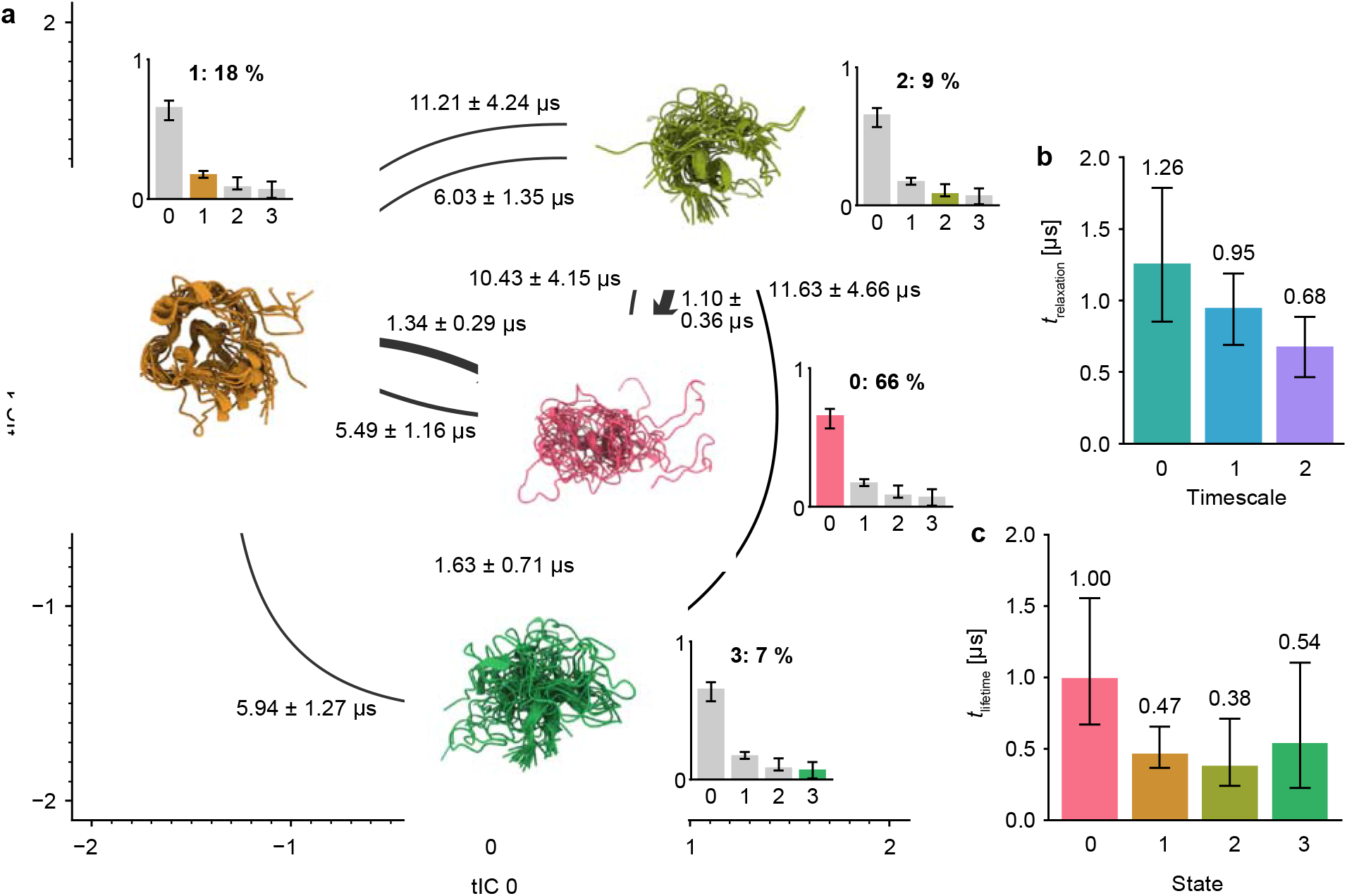
Populations and mean first-passage times in the kinetic ensemble of Aβ42. **(a)** Mean first-passage times and their standard deviations between states in the kinetic ensemble; thicker arrows correspond to higher transition probabilities. The state location is projected on to the two slowest time-independent coordinates (tICs) and the structures shown are 20 high-weight conformations from all models aligned on the most prominent secondary structure motifs (see **Figure 3a**). Transitions with mean first-passage times slower than 20 μs are not shown (**Figures S12a, c**). **(b)** Slowest relaxation timescales of the 4-state model. **(c)** Mean lifetime of each state in the 4-state model. Error bars indicate 95th percentiles of the bootstrap sample of the mean.

### Methionine-oxidized Aβ42 features faster state transitions than reduced Aβ42

To contextualize our findings and demonstrate the transferability of this approach, we also performed additional simulations of the methionine-oxidized form of Aβ42 (Aβ42-MetSO). These were carried out in three rounds of 1024 trajectories each, using the same FAST procedure reported above. We validated our model using the same approach, i.e. plotting the dependence of the relaxation timescales on the model lag time and number of trajectories (**Figure S13**) and using the Chapman-Kolmogorov test (**Figure S14**). We found that the state decomposition remains largely identical (**Figure S15**), with a population shift away from the ordered states 1, 2, and 3 towards the more extended state 0. This shift is also evident in the relaxation timescales and state transitions towards state 0, which are accelerated in the Aβ42-MetSO ensemble as compared to that of Aβ42 (**Figure S16a-b**). Additionally, the lifetime of state 0 is also vastly prolonged, from ~1 μs in the Aβ42 ensemble to ~6 μs for Aβ42-MetSO (**Figure S16c**). These findings are consistent with nuclear magnetic resonance studies on both forms, in which Aβ42-MetSO was shown to exhibit higher backbone mobility than Aβ42^55^ and overall lower β-sheet content^56^. These results coincide with higher solubility and reduced aggregation propensity^57^.

## Discussion

In this work we determined the kinetic ensemble of Aβ42 using a neural network approach. We observed that the choice of the number and quality of inputs is crucial. In particular, we observed a large increase in the VAMP2 scores upon switching from inter-residue Cα distance matrices to nearest-neighbour heavy atom distances. These results suggest that using more sophisticated and higher resolution features could further improve the state decomposition. One such method could be the use of convolutional layers^58–60^, either acting on distance matrices or on the 3D protein structure itself.

Our study also identifies the importance of developing robust mathematical tools to handle the Koopman matrix and corresponding error estimations. There have been numerous developments in the Markov model literature for these problems, such as the use of Bayesian methods to estimate both state discretization and statistical sampling uncertainties^61^ or the use of additional experimental data to improve the transition matrix estimation^62^. These tools are, however, all based on the availability of discrete transition counts, and there are no direct analogs for Koopman models so far.

Aβ42 has been studied extensively using molecular simulations in the context of thermodynamic ensembles, commonly using enhanced sampling techniques such as replica-exchange molecular dynamics or variants thereof^33–38^. These and related studies suggest that this peptide is extremely sensitive to the choice of force field and water model^63^ as well as the amount of sampling^41^. These studies used higher simulation temperatures, making a quantitative comparison with our study difficult. The work by Lin *et al.*^34^ is especially notable as the authors used a Markov state model to analyze their 200 μs simulation. They, however, did not report on the kinetic properties of the system and focused on ensemble-averaged structural features instead. Meng *et al.* also identified a large disordered population of 72 % and an end-to-end contact population of 3 % in their analysis^33^, consistent with our analysis. Rosenman *et al.* found many highly diverse clusters with populations of ~5 %, comprising both extended and locally structured conformations^35^. Sgourakis *et al.* used a spectral clustering technique based on contact maps to identify many locally structured conformations^36^.

Our results are particularly interesting when viewed in context of the kinetic hub model of protein folding^64^. In this model, the native state of a folded protein is the most populated state, quickly reachable from other partially or completely unfolded states. This model can be seen as the kinetic complement of the thermodynamic funnel concept of protein folding^65^. We may therefore propose an analogous model for disordered proteins, i.e. a kinetic counterpart to the recently proposed inverted funnel concept^38^. Aβ42 clearly adopts a highly populated disordered state, with slow transitions into less populous, partially folded states. We can thus view this kinetic geometry as an ‘inverted kinetic hub’.

Additionally, the ability to capture such dramatic structural and dynamic differences between the Aβ42 and Aβ42-MetSO peptides, which only differ by a single atom, in a way that is consistent with independent experimental data demonstrates the power of the soft Markov state model method for studying the complexity of disordered proteins.

## Conclusions

We have presented a general strategy to characterize the structure, thermodynamics, and kinetics of disordered proteins through kinetic ensembles. To illustrate the application of this approach, we generated two kinetic ensembles of Aβ42, a disordered peptide closely associated with Alzheimer’s disease, including the reduced and oxidised forms of Met35. Our results show that the use of soft Markov state models enables one to find a suitable state-space discretization to generate kinetic ensembles, a problem that has been challenging to date, and offers insight into the structure, dynamics, and function of these complex biomolecules. Our findings also suggest that soft Markov models and clustering approaches based on neural networks could be applicable to other systems characterized by fast state transitions and conformational heterogeneity.

## Methods

### Simulation details

All simulations were carried out with GROMACS 2018.1^66^ using 1,024 individual Virtual Machine instances on Google Compute Engine. All instances used the compute-optimized n1-highcpu-8 machine type, each configured with 8 Intel Haswell CPU cores, 7.2 GB of RAM, and 100 GB of disk space. 1,024 Aβ42 starting conformations were chosen from a previous metadynamics-biased molecular dynamics simulation by sample-weighted *k*-Means clustering on the space of backbone dihedral angles and picking a random conformation from each of the 1024 clusters. Each conformation was solvated separately in a rhombic dodecahedron box with a volume of 358 nm^3^ using between 11,711 and 11,751 water molecules. The system was minimized using the steepest descent algorithm to a target force of less than 1000 kJ / (mol / nm). Equilibration was performed over a time range of 500 ps in the NVT ensemble using the Bussi thermostat^67^ and 500 ps in the NPT ensemble using Berendsen pressure coupling^68^ while applying a position restraint on all heavy atoms. Production simulations were carried out at 278 K using the CHARMM22* force field^69^ and the TIP3P water model^70^ using a 2 fs timestep in the NVT ensemble. Electrostatic interactions were modelled using the Particle-Mesh-Ewald approach^71^ with a cut-off for the short-range interactions of 1.2 nm. Constraints were applied on all bonds with the LINCS algorithm^72^ using a matrix expansion on the order of 4 and 1 iteration per step. Simulations were carried out with the initial 1,024 starting conformations, the resulting trajectories were then used with the fluctuation amplification of specific traits (FAST) approach^39^ to choose 1,024 new starting structures. The clustering for FAST was conducted by first performing time-lagged independent component analysis (TICA)^15,16^ with a lag time of 5 ns using Cα-distance matrices as input and the *k*-Means algorithm in this reduced space to create 128 clusters. We chose to both maximize the deviation to the mean Cα-distance matrix for each cluster and maximize the sampling of existing clusters using a balance parameter of α = 1.0. All amino acids were weighted equally. This procedure was repeated three times to yield a total of 5,119 trajectories, with an aggregated simulation time of 315 μs. The shortest and longest trajectory lengths were 9.75 and 90.5 ns respectively. The trajectories were subsampled to 250 ps time steps to yield 1,259,172 frames.

Simulations of methionine-oxidized Aβ42 were performed analogously in three rounds using the same FAST scheme and simulation parameters yielding 3071 trajectories with an aggregated simulation time of 317 μs and 1,268,139 frames. Oxidized methionine parameters were constructed based on the parameters for dimethyl sulfoxide from the CHARMM generalized force field^73^. Initial conformations were created by sampling 1024 structures from the Aβ42 ensemble and mutating the methionine residues to the oxidized form.

### Neural network

State decomposition was performed using the VAMPNet approach with physical constraints.^21,22^ The two-lobed neural network takes as input a pair of frames separated by some lag time and yields a state assignment vector as output. We chose inter-residue nearest-neighbour heavy atom distance matrices with first- and second-degree off-diagonals removed as input (780 dimensions) and used between 2 and 8 output nodes with softmax activation. The architecture of the χ layers of the neural network follows the self-normalizing setup described by Klambauer et al.^47^, using scaled-exponential linear units, normal LeCun weight initialization, and alpha dropout. The constrained part is composed of two additional layers **u** and **s** which are trained with and without the *χ* layers, successively. The VAMP2 loss function for the *χ*(**x**_*t*_) network is given as follows:

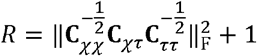

where

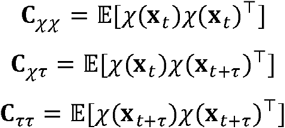

The constrained network is trained using the VAMP-e score^22^ *R*:

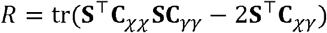

where

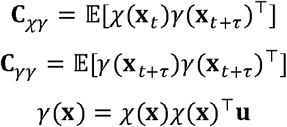

and **s** and **u** are the trainable reversibility and non-negativity constraints, respectively. Error estimates were obtained through bootstrap aggregation (bagging), i.e. by training 20 independent neural networks with independently randomized and shuffled 9:1 train-validation splits, and, to prevent overfitting, stopping training when the validation score no longer improved. The model was implemented using Keras 2.2.4^74^ with the Tensorflow 2.1.0^75^ backend. The neural network parameters were chosen through two successive random grid searches with scikit-optimize 0.5.2^76^, first using a coarse grid spanning a large parameter space, then a finer grid over a local space around the optimum parameters. We found the best parameters to be a network lag time of 5 ns, a layer width of 256 nodes, a depth of 5 layers, an *L*_2_ regularization strength of 10^−8^ and no dropout. Training was performed on a single Google Compute Engine instance with an NVidia Tesla V100 GPU, 12 Haswell cores, and 78 GB of RAM. Training of the *χ* model was performed using batch sizes of 10000 frame pairs and the Adam minimizer with a learning rate of 0.05, β_2_ = 0.99 and 0.0001. Overfitting was addressed through early stopping, i.e. training was stopped when the VAMP validation set score did not increase by at least 0.001 over the previous 5 epochs. The constraint layers **u** and **s** as well as the full network including the *χ* layers were trained with the Adam minimizer with a learning rate of 0.0005. The constraint layers were trained with batches the size of the full dataset, with early stopping with no improvement in the loss after 10 epochs.

### Analysis

The output of a single neural network is a state assignment vector *χ*(**x**_*t*_) of each frame **x**_*t*_ of the simulation. The ensemble averaged value of an observable *A*(**x**_*t*_) for a state *i* is therefore an average weighted by the state assignment for *T* time steps:

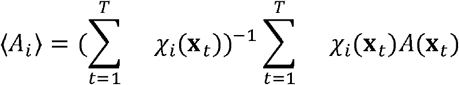

Ensemble averaged quantities can be calculated by first computing a weight *w*_*t*_ for each frame of the simulation based on the state assignment *χ*(**x**_*t*_) at time *t* and the equilibrium distribution *π*:

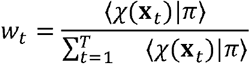

The ensemble averaged observable ⟨*A*⟩ can then be calculated as the weighted average:

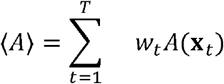

To estimate the error, we sort each state assignment vector, as multiple trained neural networks do not necessarily conserve the order of identified states. To do so, we calculate the mean inter-residue nearest-neighbour heavy atom distance matrix for each identified state and sort the states based on the lowest root-mean-square deviation between these matrices. We also ensure that the sorting is unique, i.e. we are never duplicating state assignments. The Koopman matrix **K**(*τ*) is then estimated directly from the constrained neural network.

Internal model validation is performed with the Chapman-Kolmogorov test (**Figures S5-7, and S14**):

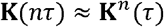

i.e. we expect a model estimated at some lag time *τ* to behave the same way as one estimated at a multiple *nτ* of it. The correct lag time *τ* for the model is estimated by plotting the relaxation (also known as implied) timescales *t*_*i*_ (**Figures 4a and S13a**):

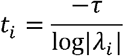

where *λ*_*i*_ is the *i*th eigenvalue of the Koopman matrix **K**(*τ*):

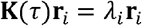

As the Koopman matrix is row-stochastic, the largest eigenvalue is 1, and its associated eigenvector is the stationary distribution *π* of the system. The state lifetimes 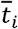 are given by

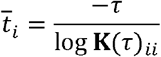

where **K**(*τ*)_*ii*_ are the diagonal entries of the Koopman matrix.

Chemical shifts were back calculated with CamShift^48^ as implemented in PLUMED 2.4.1^77,78^ using the full trajectories, averaged as described above and compared to previously recorded NMR data^31^. Time-lagged independent component analysis was performed with inter-residue nearest-neighbour heavy atom distances as input, using a lag time of 5 ns and using kinetic maps^79^.

### Conventional Markov state model

Markov state models based on discrete states were constructed as follows: Time-lagged independent component analysis was performed with inter-residue nearest-neighbour heavy atom distances as input, with a lag time of 5 ns. Various clustering methodologies were then evaluated using the 5-fold cross-validated generalized matrix Rayleigh quotient score using every 50th frame of the full trajectory (25183 frames) and a lag time of 12.5 ns^43^. Evaluation was performed over the number of time-lagged independent components, clusters, and algorithm (minibatch *k*-Means^80,81^, minibatch k-Medoids^82^, Gaussian mixture model) using MSMBuilder^83^ (**Figure S1**). 100 Markov models with the highest scoring parameters were then sampled and the timescales evaluated for the conventional Bayesian (**Figure S2a**)^12,84^, hidden Markov model (**Figures S2b-d**)^44^ and Perron cluster-cluster analysis^45^ (**Figures S2e-f**) cases.

### Errors

Observable errors for each state were calculated by taking a trajectory sample based on the training data for a neural network sample. All statistics were then calculated on the bootstrap sample, and the errors reported in the figures show the 95th percentile of this sample unless noted otherwise.

## Code and data availability

Subsampled trajectory and intermediate data, as well as the trained neural network weights and analysis notebooks are available from https://zenodo.org/record/4050508.

## Acknowledgements

GTH is supported by the Rosalind Franklin Research Fellowship at Newnham College. The authors would like to thank Dr. Steven Kearnes for useful feedback and suggestions for the manuscript and Dr. Massimilliano Bonomi for useful discussions and suggestions. The authors also acknowledge support from Google and the Google Accelerated Science team for providing access to the Google Cloud Platform for simulations and analysis.

