## Supplementary Information for "A kinetic ensemble of the Alzheimer’s Aβ peptide"

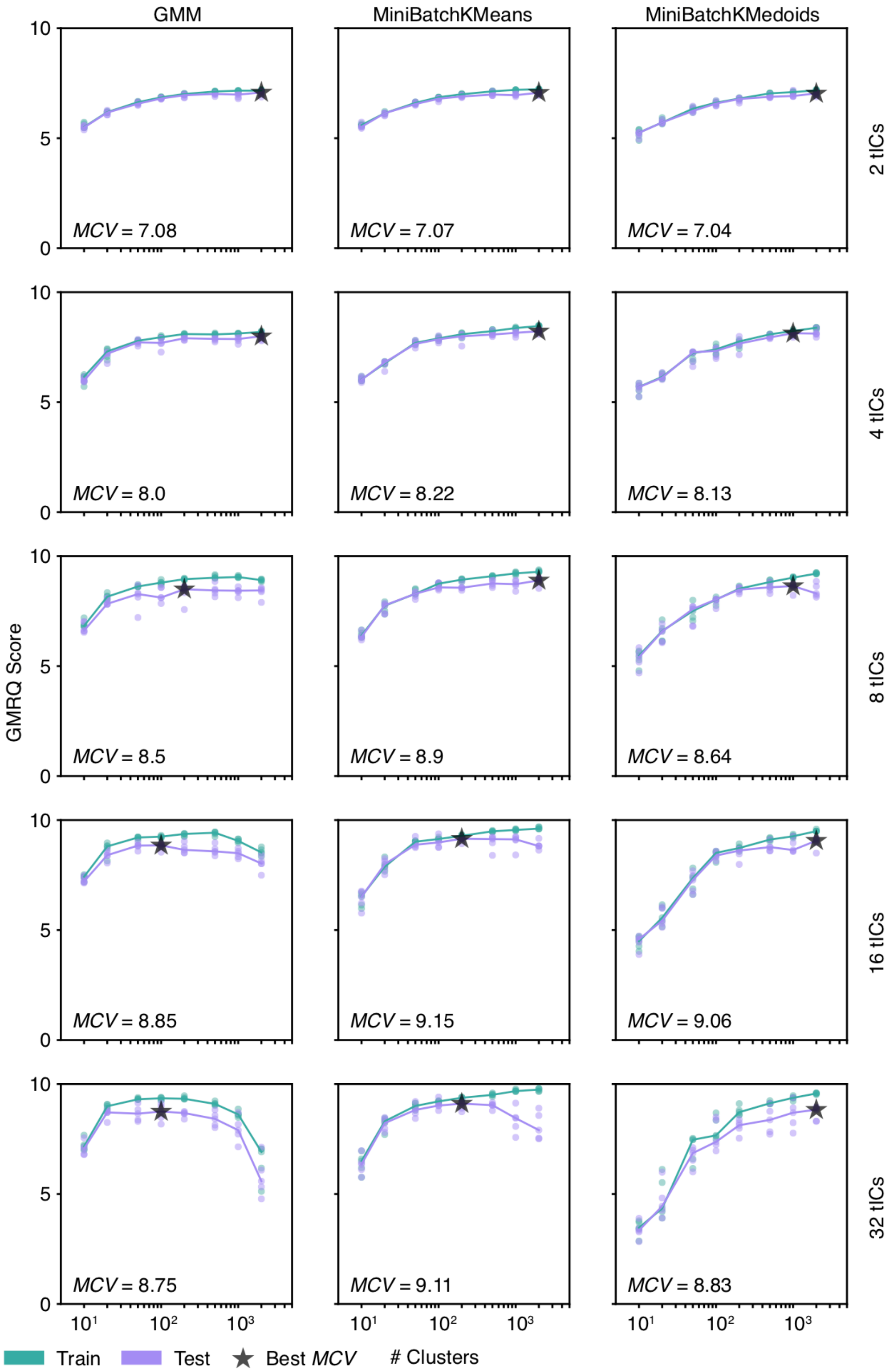


**Figure S1.** Hyperparameter scan for a conventional discrete-state Markov state model. Clustering algorithms evaluated include Gaussian mixture models, minibatch k-Means, and minibatch k-Medoids using between 10 and 2000 microstates and between 2 and 32 input dimensions. The mean cross-validation score (MCV) is shown in the bottom left.

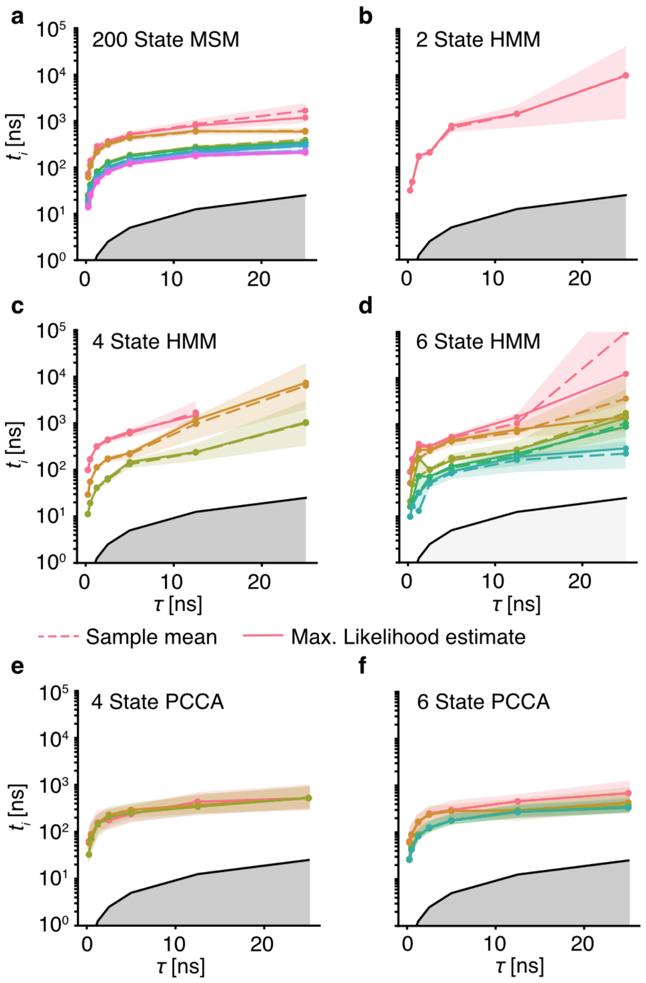


**Figure S2.** Relaxation (implied) timescales for conventional discrete-state Markov state models, showing inability to construct a model with converging timescales. **(a)** Relaxation timescales for a 200-microstate model as a function of model lag time. **(b-d)** Relaxation timescales for hidden Markov state models using 2, 4, and 6 output states respectively, as a function of model lag time. **(e-f)** Relaxation timescales for 4 and 6-state Markov state models built using Perron cluster-cluster analysis (PCCA) from the 200-microstate model as a function of model lag time. Shaded areas indicate 95% confidence intervals of the sample mean.

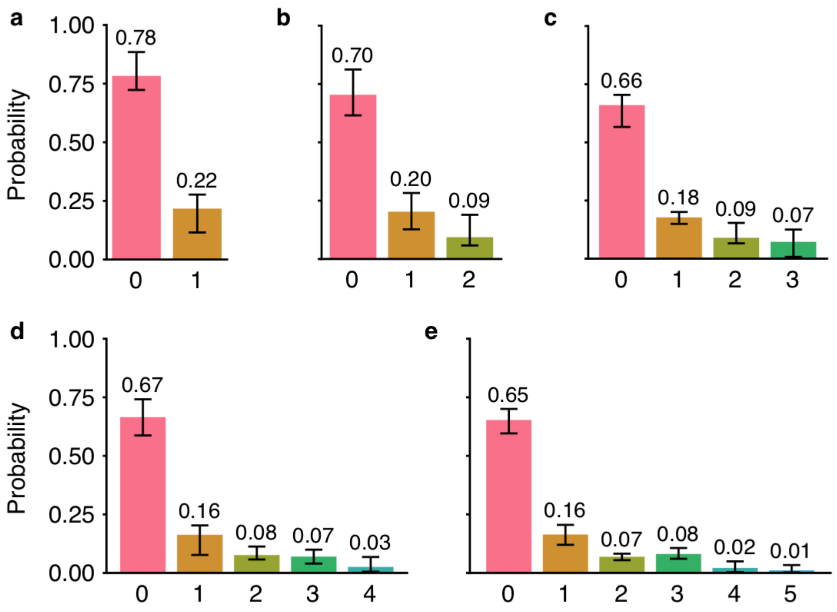


**Figure S3.** Equilibrium distribution of the **(a)** 2, **(b)** 3, **(c)** 4, **(d)** 5, and **(e)** 6-state models. Error bars indicate 95th percentiles of the model mean.

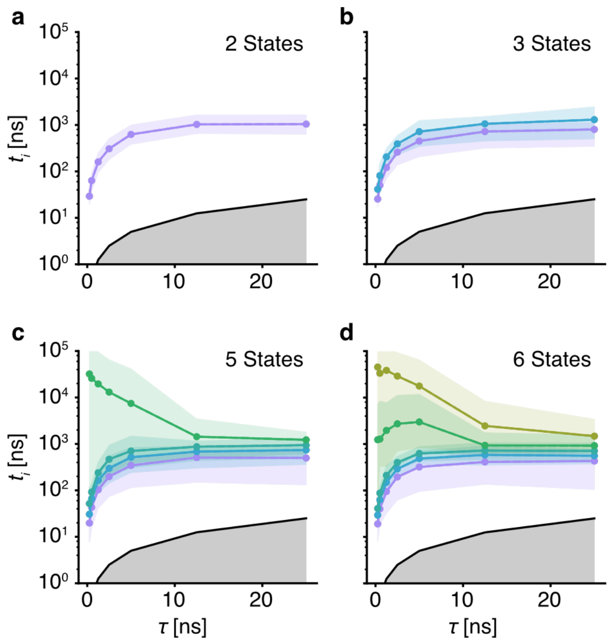


**Figure S4.** Relaxation (implied) timescales of the **(a)** 2, **(b)** 3, **(c)** 5, and **(d)** 6-state models as a function of model lag time. Shaded areas indicate 95th percentiles of the model mean.

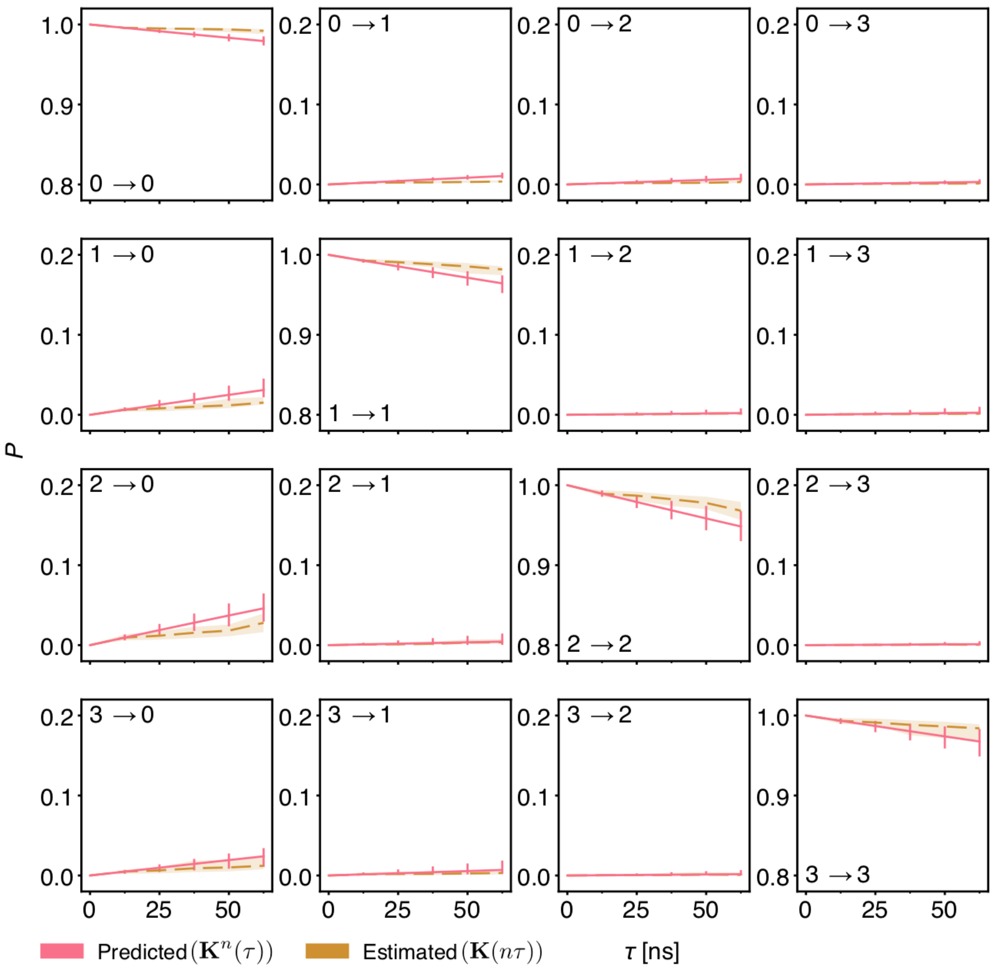


**Figure S5.** Chapman-Kolmogorov test for the 4-state model. Each panel indicates the transition probability for one matrix entry for successive applications and estimations of the Koopman matrix. Shaded areas and error bars indicate 95th percentiles of the model mean.

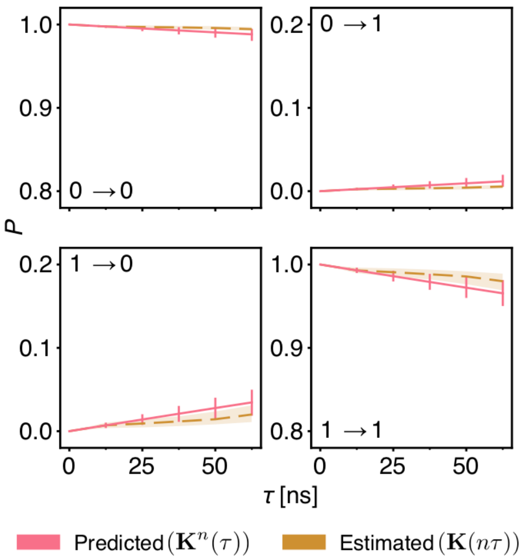


**Figure S6.** Chapman-Kolmogorov test for the 2-state model. Each panel indicates the transition probability for one matrix entry for successive applications and estimations of the Koopman matrix. Shaded areas and error bars indicate 95th percentiles of the model mean.

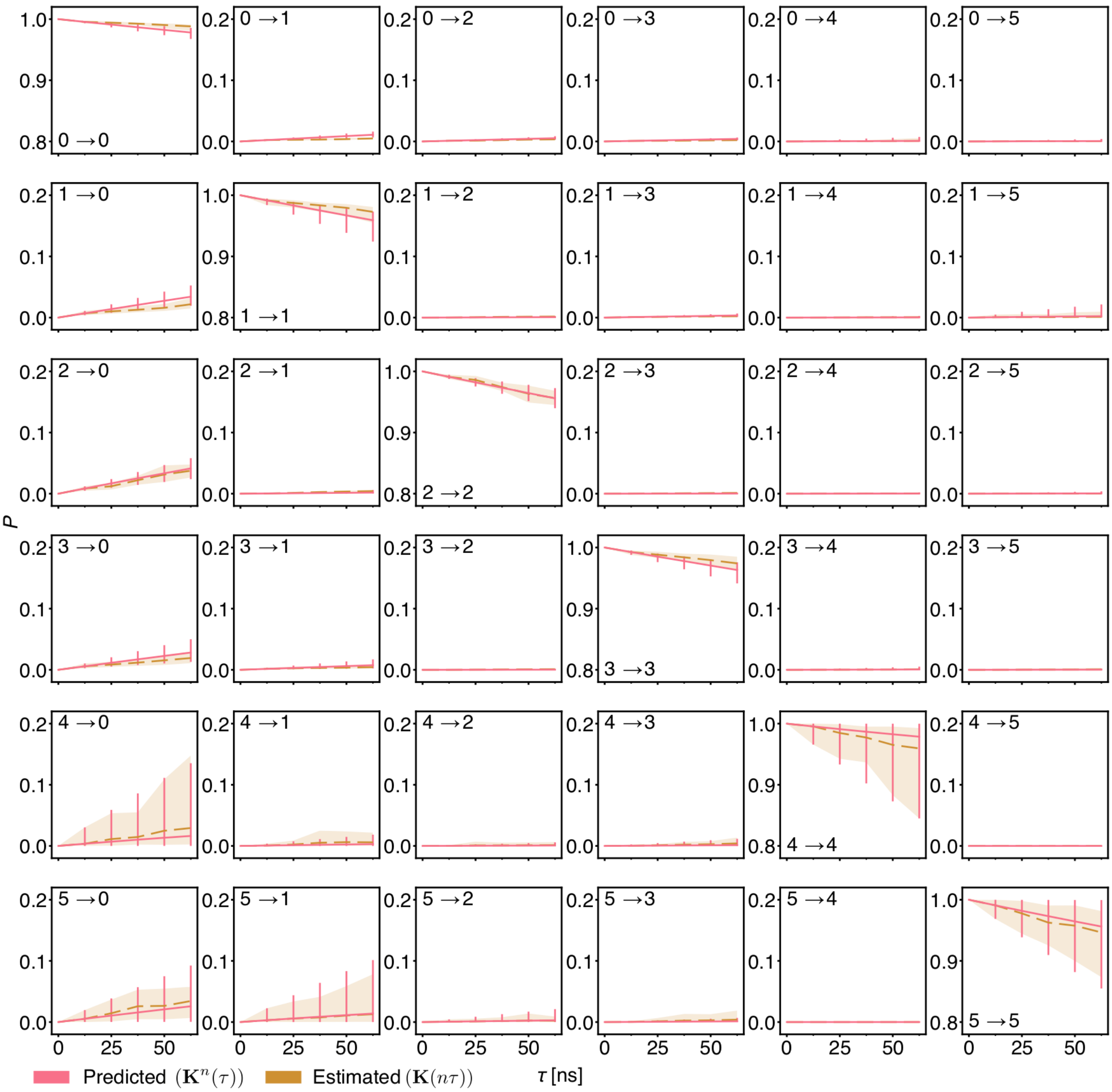


**Figure S7.** Chapman-Kolmogorov test for the 6-state model. Each panel indicates the transition probability for one matrix entry for successive applications and estimations of the Koopman matrix. Shaded areas and error bars indicate 95th percentiles of the model mean.

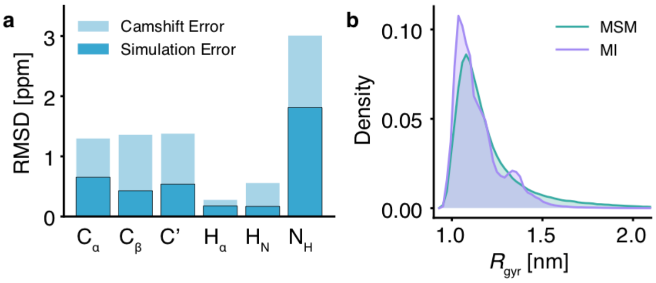


**Figure S8. (a)** Root-mean-square deviations between experimentally determined NMR chemical shifts and those back calculated using CamShift[1]; the deviations are smaller than the intrinsic CamShift errors. **(b)** Comparison of the probability distributions of the radius of gyration computed for the current Markov state model (MSM, green) and the previously performed metadynamic metainference simulations (MI, purple)[2].

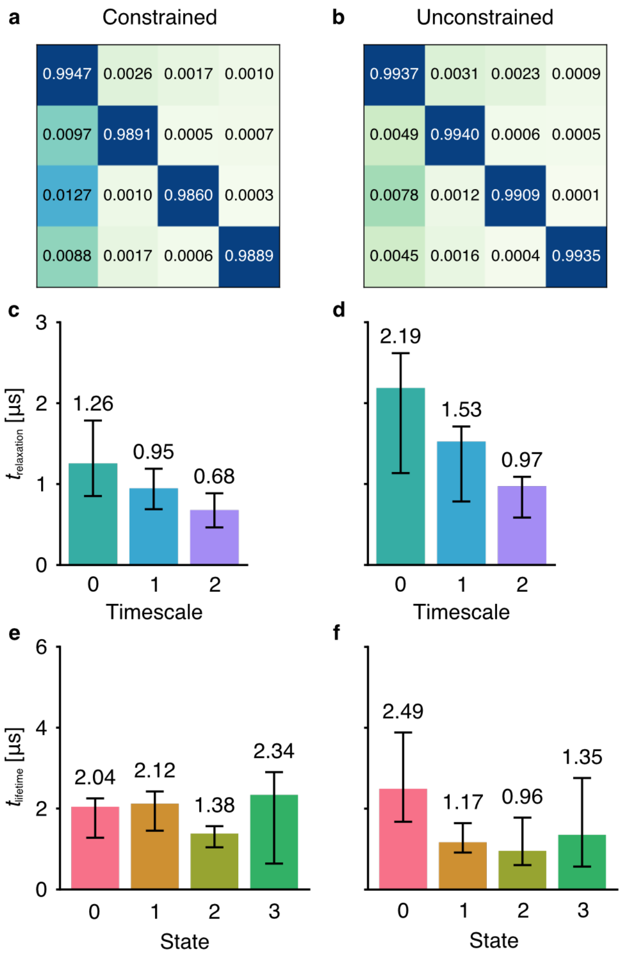


**Figure S9.** Comparison of the 4-state model constrained to feature reversibility and positive transition matrix elements and the unconstrained 4-state model. **(a-b)** Mean transition matrix elements (state transition probabilities). **(c-d)** Relaxation timescales of the constrained and unconstrained 4-state models. **(e-f)** State lifetimes for the constrained and unconstrained models. Error bars indicate 95th percentiles of the model mean.

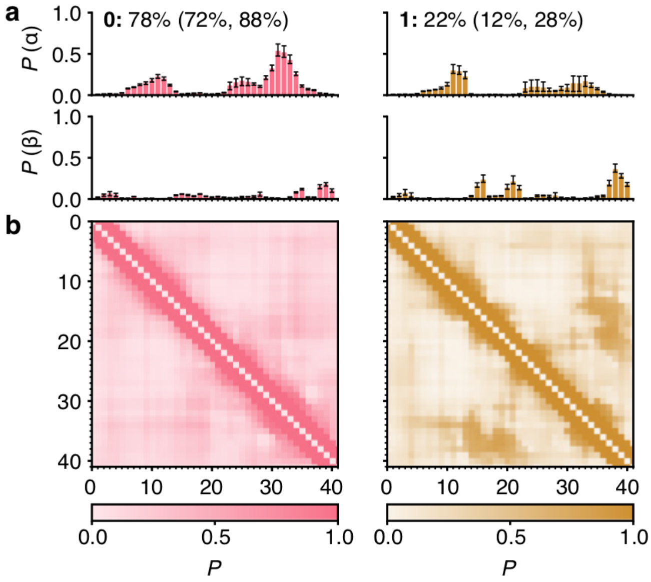


**Figure S10.** Structural properties of Aβ42 in the two-state model. **(a)** Populations of α-helical and β-sheet content per residue as calculated using DSSP[3]. The equilibrium percentage of each state is given above, with the 95th percentile in parentheses. **(b)** Contact probability maps with a cut-off of 0.8 nm. Error bars indicate 95th percentiles of the bootstrap sample of the mean.

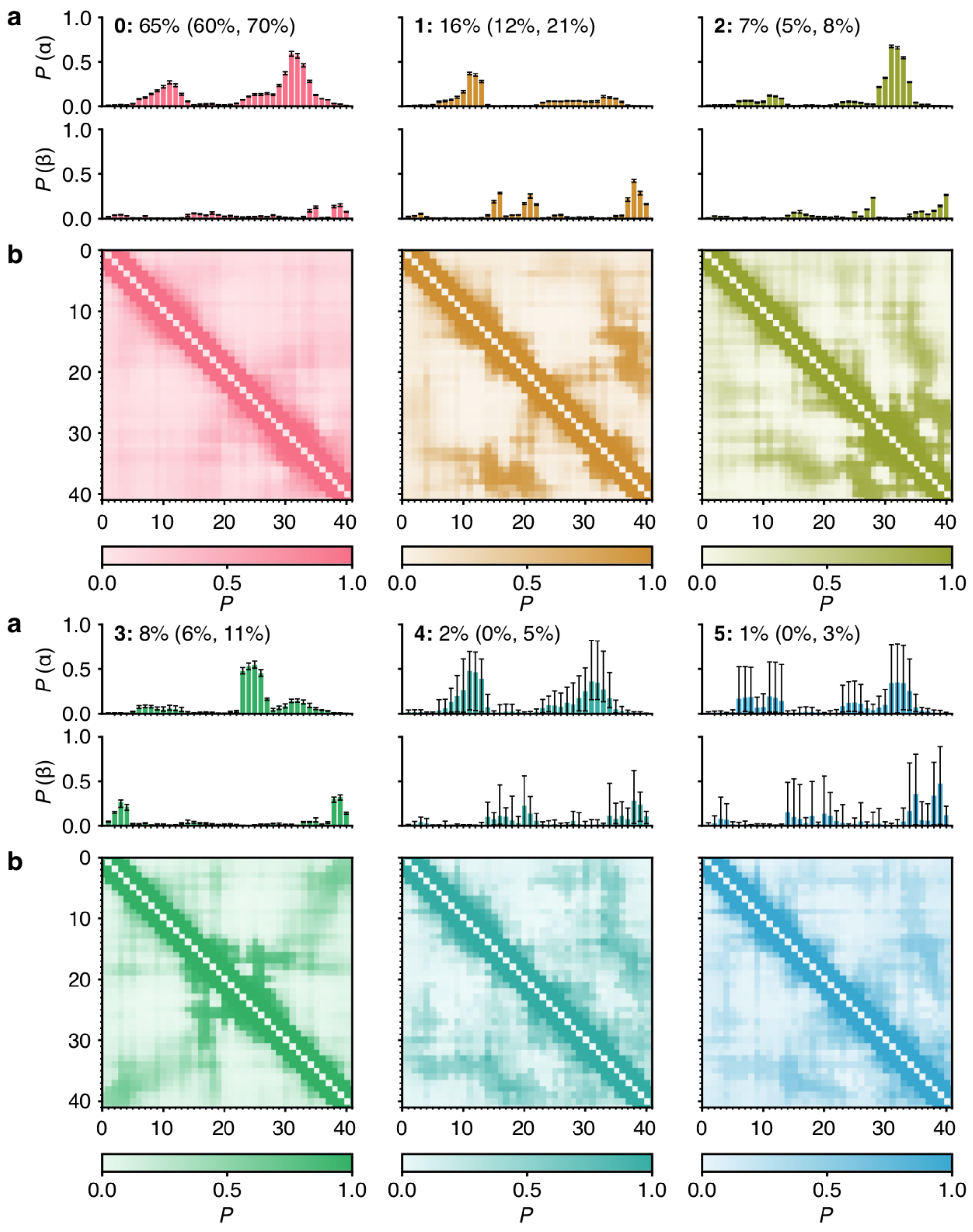


**Figure S11.** Structural properties of Aβ42 in the six-state model. **(a)** Populations of α-helical and β-sheet content per residue as calculated using DSSP[3]. The equilibrium percentage of each state is given above, with the 95th percentile in parentheses. **(b)** Contact probability maps with a cut-off of 0.8 nm. Error bars indicate 95th percentiles of the bootstrap sample of the mean.

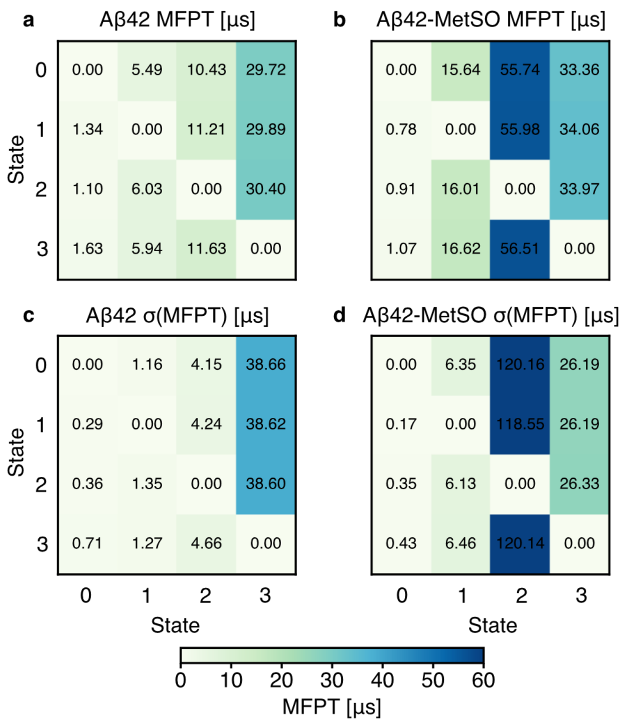


**Figure S12.** Full mean first-passage times for **(a)** Aβ42 and **(b)** Aβ42-MetSO in μs. **(c-d)** Standard deviations for the mean first-passage times of both models in μs, across all models.

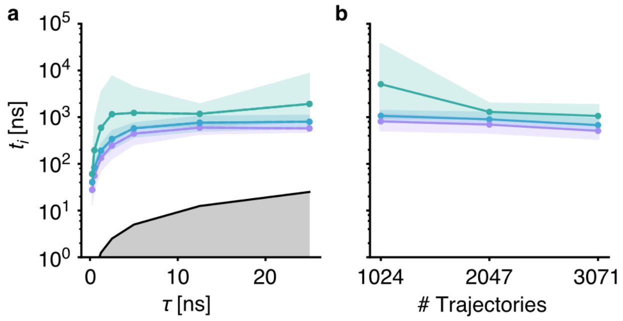


**Figure S13.** Computational validation of the Aβ42-MetSO kinetic ensemble. **(a)** Dependence of the three longest relaxation timescales (green, cyan and purple, respectively) on the lag time τ. The grey shading indicates the timescales for which the Koopman model can no longer resolve the relaxation timescales. **(b)** Dependence of the relaxation timescales on the number of trajectories used to build the kinetic ensemble as a 4-state model. Shaded areas indicate 95th percentiles of the bootstrap sample of the mean.

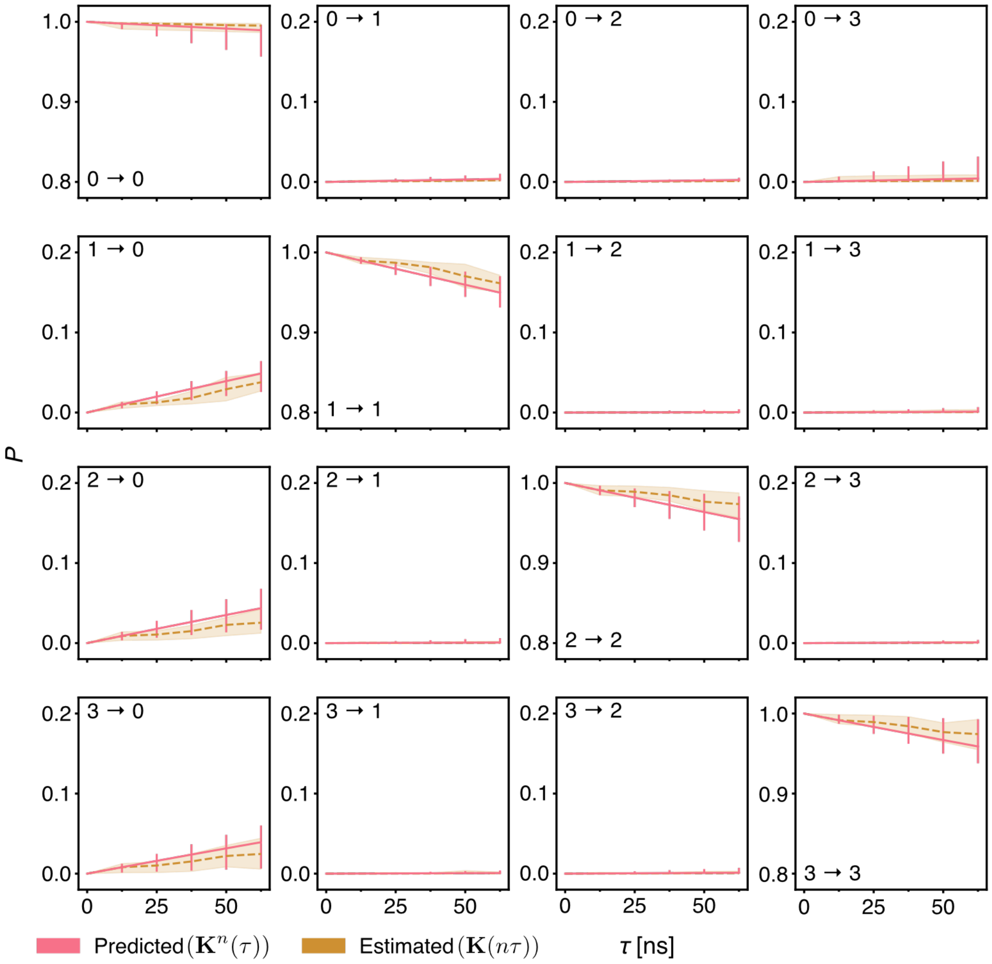


**Figure S14.** Chapman-Kolmogorov test for the 4-state model of Aβ42-MetSO. Each panel indicates the transition probability for one matrix entry for successive applications and estimations of the Koopman matrix. Shaded areas and error bars indicate 95th percentiles of the model mean.

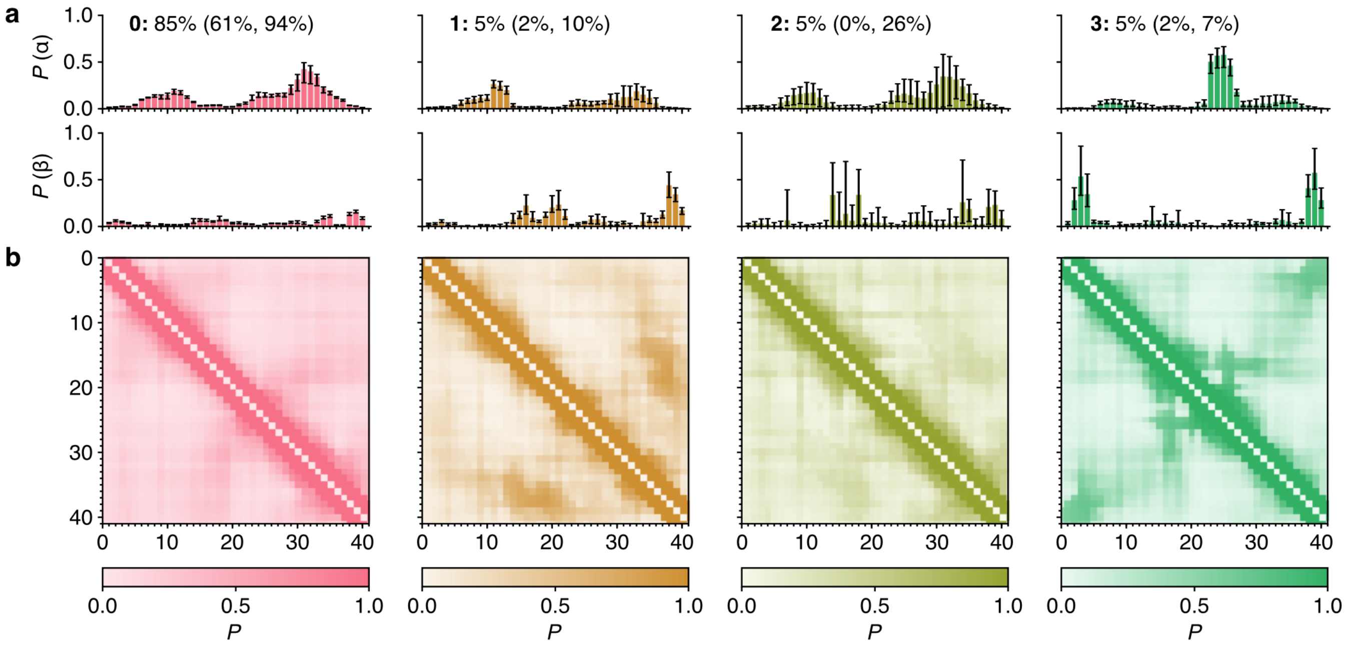


**Figure S15.** Structural properties of Aβ42-MetSO in the four-state model. **(a)** Populations of α-helical and β-sheet content per residue as calculated using DSSP[3]. The equilibrium percentage of each state is given above, with the 95th percentile in parentheses. **(b)** Contact probability maps with a cut-off of 0.8 nm. Error bars indicate 95th percentiles of the bootstrap sample of the mean.

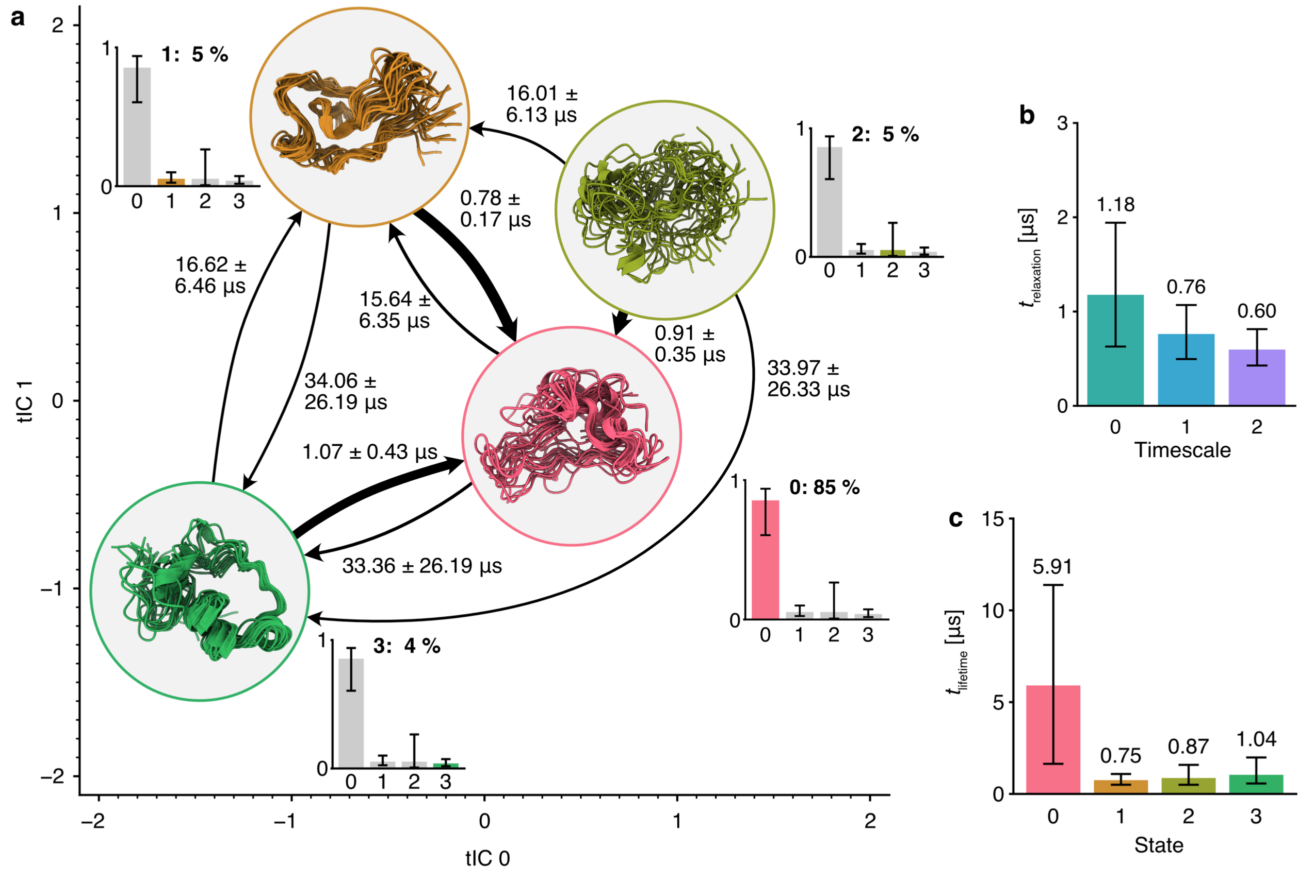


**Figure S16.** Populations and mean first-passage times in the kinetic ensemble of Aβ42-MetSO. **(a)** Mean first-passage times and their standard deviations between states in the kinetic ensemble; thicker arrows correspond to faster transitions. The state location is projected on to the two slowest time-independent coordinates (tICs) and the structures shown are 20 high-weight conformations from all models aligned on the most prominent secondary structure motifs (see **Figure 3a**). Transitions with mean first-passage times slower than 40 µs are not shown (**Figures S16b, d**). **(b)** Slowest relaxation timescales of the 4-state model. **(c)** Mean lifetime of each state in the 4-state model. Error bars indicate 95th percentiles of the bootstrap sample of the mean.
